# The impact of population variation in the analysis of microRNA target sites

**DOI:** 10.1101/613380

**Authors:** Mohab Helmy, Andrea Hatlen, Antonio Marco

**Affiliations:** School of Biological Sciences, University of Essex, Colchester CO4 3SQ, United Kingdom; Faculty of Life Sciences & Medicine, King’s College London, London SE1 9RT, United Kingdom

**Keywords:** miRNAs, human populations, gene regulation, microRNA target prediction, evolution

## Abstract

The impact of population variation in the analysis of regulatory interactions is an underdeveloped area. MicroRNA target recognition occurs via pairwise complementarity. Consequently, a number of computational prediction tools have been developed to identify potential target sites, that can be further validated experimentally. However, as microRNA target predictions are done mostly considering a reference genome sequence, target sites showing variation among populations are neglected. Here we study variation at microRNA target sites in human populations and quantify their impact in microRNA target prediction. We found that African populations carry a significant number of potential microRNA target sites that are not detectable in the current human reference genome sequence. Some of these targets are conserved in primates and only lost in Out-of-Africa populations. Indeed, we identified experimentally validated microRNA/transcript interactions that are not detected in standard microRNA target prediction programs, yet they have segregating target alleles abundant in non-European populations. In conclusion, here we show that ignoring population diversity may leave out regulatory elements essential to understand disease and gene expression, particularly neglecting populations of African origin.

## Introduction

The study of gene regulatory iterations are at the heart of biomedical research. Many disease-causing mutations affect regulatory motifs that eventually lead to a miss-regulation of fine-tuned biological processes. In the last years, microRNAs have emerged as an important type of regulatory molecules involved in virtually all biological networks [1,2]. Both the deletion or overexpression of microRNAs have been associated to human diseases (see for instance the compilation by [3]). Notably, the role of microRNAs in cancer is a particularly active research field [4,5]. Unlike other regulatory molecules (such as transcription factors or RNA binding proteins) the interaction between regulator and target is mediated by pairwise nucleotide complementary [1]. Specifically, a microRNA is partially complementary to motifs in RNA transcripts and, after binding, transnational repression or even RNA degradation mechanisms are triggered [1,6]. Indeed, changes at microRNA target sites are now associated to major human diseases including cancer, neurodegeneration and metabolic disorders (see [7,8] and references therein), among others. The study of microRNA target sites is becoming more important as we start to understand the complexities behind microRNA regulatory networks.

From the computational point of view, the microRNA targeting mechanism have a clear advantage: targets can be predicted by scanning genomic sequences for potential complementary sites [9–11]. However, microRNA target prediction algorithms have, in general, high rates of false positives [12,13]. Consequently, most research on microRNA targets complements a computational sieve of potential target sites plus experimental validations, usually by luciferase assays or similar (e.g. [14]). Therefore, the first step in microRNA target analysis is the prediction of binding sites from a primary sequence. In biomedical research, this primary sequence is often, if not always, the human reference genome sequence (currently hg38). However, a reference genome does not take into account existing variation within human populations and, therefore, any inference from such a sequence may be biased. Populations of African origin show a higher nucleotide diversity than other populations, probably as a consequence of a recent origin of Eurasian populations from a small group of African migrants (reviewed in [15]). There exist several projects aimed to account for all this nucleotide variability in human populations (i.e. [16]). Nevertheless, microRNA target prediction programs rely on reference genome sequences (as other regulatory interaction programs do), where this diversity is not truly represented.

Variation at gene regulatory sites, particularly at transcription factor binding sites, has been studied in the past (e.g. [17,18]). In microRNA target sites, some studies have shown that, as expected, functional target sites are often conserved in populations [19,20]. On the other hand, the creation of novel, potentially harmful, microRNA target sites, is selected against in populations [21,22]. In summary, the literature clearly shows evidence that nucleotide variation occurs at microRNA target sites. The question is, how much of this variation is neglected in the human reference genome sequence, and therefore not accounted in standard microRNA target prediction programs? In other words, what is the impact of nucleotide variation among humans in microRNA target prediction research? This is the question we are tackling in this paper.

## Results and Discussion

We compiled biallelic SNPs at predicted microRNA target sites (see Methods) for which one of the alleles is a target site and the other is not a target site. These are the target/near-target pairs as defined before [11,21]. Considering that populations of African origin host more variation, including ancestral alleles, than other populations, we hypothesize that ancestral target sites that are not present in the reference genome sequence were lost in Out-of-Africa populations. Thus, we first consider target sites whose allele in the reference genome sequence is not a target yet the ancestral allele was a target site. The distribution of these alleles frequencies in five major groups (African, European, South Asian, Native American and East Asian) indicates that non-African populations tend to have the non-target allele whilst African populations show a wider variation, and a paucity of non-targets compared to the other populations (Figure 1, top). Additionally, we consider target sites in the reference genome sequence that were not ancestral to human populations and, as expected, we found that African populations have lower frequencies of the target allele (Figure 1, bottom). These results suggest that a significant loss and gain of microRNA target sites happened in Out-of-African populations.

**Figure 1.**
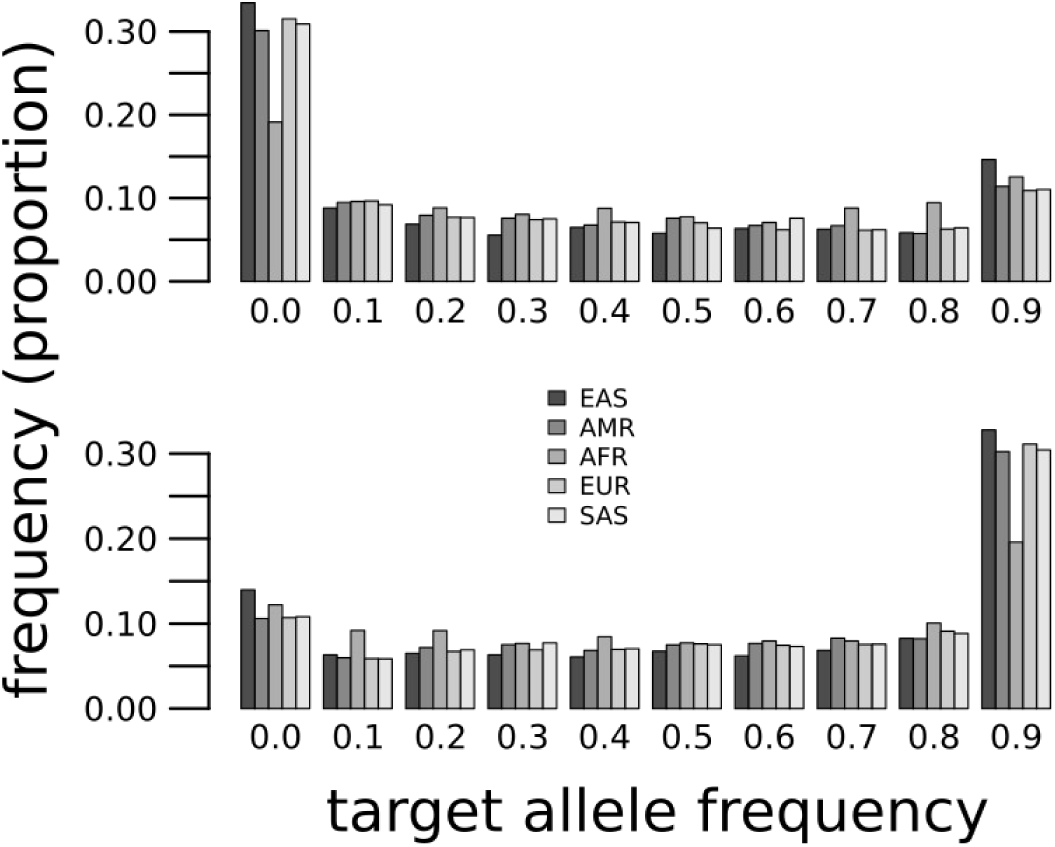
Target allele frequencies at microRNA target sites where the target is ancestral and not in the reference genome sequence (top) or the target is derived and present in the reference genome sequence (bottom).

To illustrate the impact of variation at target sites we selected SNPs with a very high degree of population differentiation (Fst > 0.7; see Methods). We found 11 target sites with such a degree of variation across populations (Table 1) for highly expressed microRNAs. In general, these target sites are lost in Out-of-Africa populations, suggesting that losses rather than the gains of target sites are more likely to happen during differentiation between populations (see discussion in [22]).

**Table 1.**
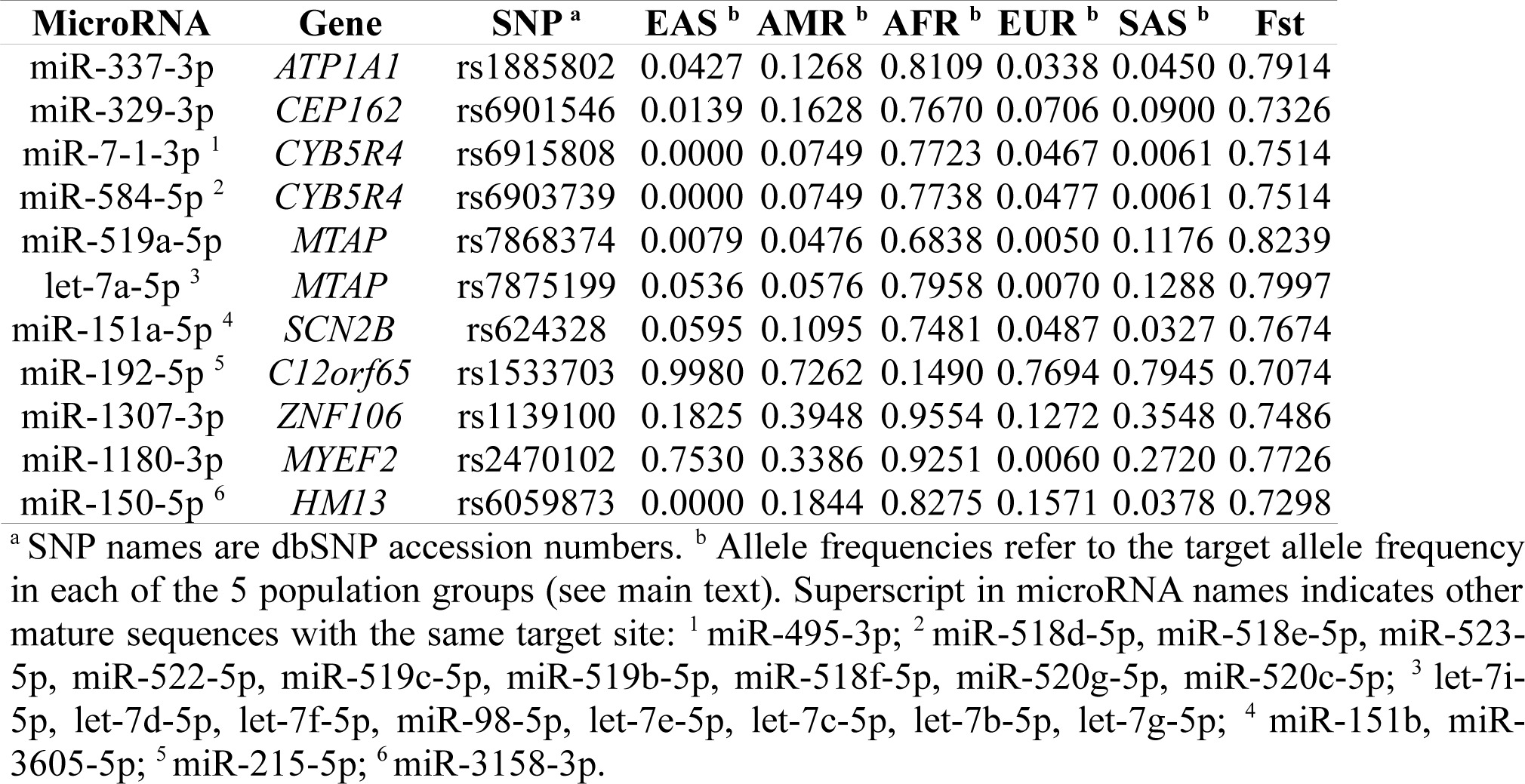
MicroRNA target sites with a very high differentiation among human populations.

We observed that *MTAP* have lost two target sites for two highly expressed microRNAs in non-African populations. These two target sites were the ancestral form in human populations and conserved as targets in other primates (Figure 2a). Genome-wide association studies have linked variants at *MTAP* with the abundance of naevi (moles) and also with melanoma incidence [23,24], although the association between *MTAP* and melanoma seems to be rather complex [23]. We speculate that the loss of two target sites at MTAP transcripts could be associated with an elevated expression of the gene in Out-of-Africa populations, perhaps by a relaxation on the selective pressures that maintained the target sites under purifying selection in African populations, where light exposure is higher, and an overproduction of MTAP may be linked to a higher incidence of skin cancer. Under this hypothesis, we expect the target allele frequency to be also negatively correlated with the latitude of sampling of non-African human populations. We compiled sampling information for human populations (see Methods) and build a linear model to predict the target allele frequency as a function of the latitude. We observed that populations from higher latitudes have lower target allele frequencies at both of the studied target sites (Figure 2b), although the statistical support was weak (rs7875199: R^2^(adjusted) = 0.18, p = 0.06; rs7868374: R^2^(adjusted) = 0.14, p = 0.09).

**Figure 2.**
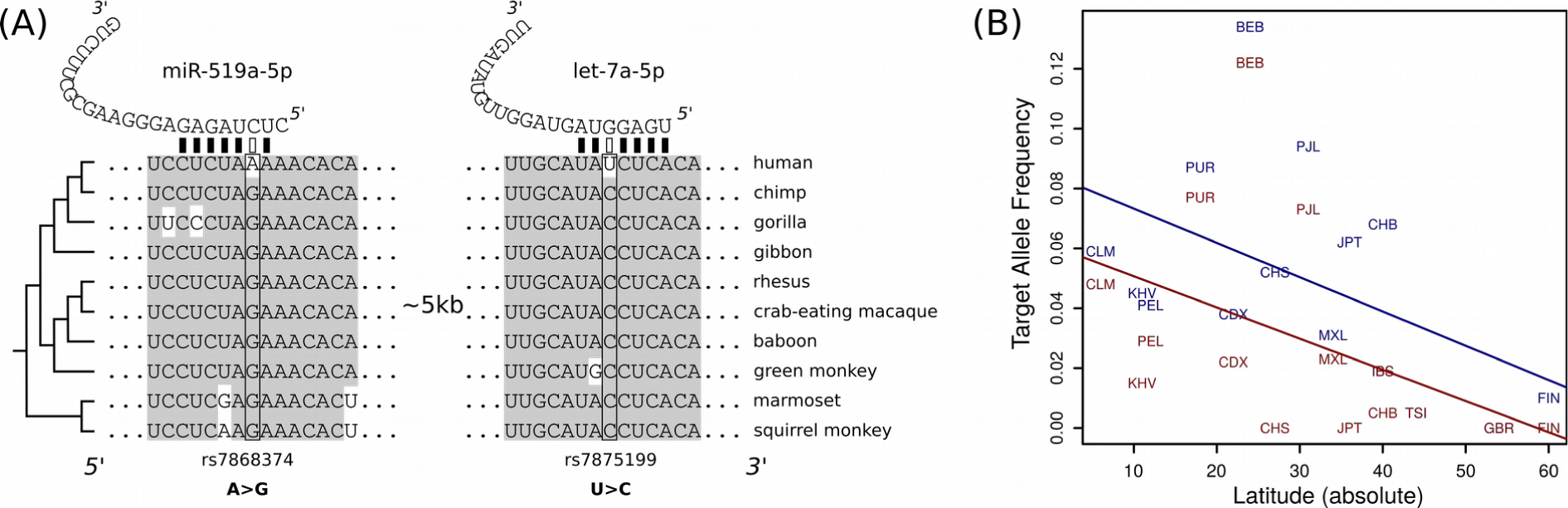
MicroRNA target sites variation at transcript from the MTP gene: (**a**) Alignment of 3’UTR fragments from 10 primate species, including human, and the location of the target sites and the polymorphic nucleotides; (**b**) Scatter-plot between the latitude (in absolute value) and the target allele frequency for target sites for miR-519a-5p (red) and let-7a-5p (blue). Straight lines are fitted linear models (see main text).

Likewise, we found a similar pattern in two target sites at *CYB5R4*, a gene associated to gene-environment interactions [25,26]. In particular, *CYB5R4* has been associated to the risk of diabetes, and variants at its 5’ region have been found to be specific of African populations [25]. The association between population variants and disease-related genes such as *MTAP* and *CYB5R4* remains highly speculative, yet it is very suggestive. In any case, the identified target sites at *MTAP* and *CYB5R4* are not present in the human genome reference genome yet they are highly abundant in African populations and conserved in primates, indicating that target prediction programs often neglect an important part of human variability.

Last, we identified microRNA/gene interactions that have been validated experimentally with a significant population differentiation (see Methods). Of the eight targets identified (Table 2), half of them were not detected as targets in the reference genome sequence yet they were identified as polymorphic. For instance, miR-338-3p has been reported to target *MACC1*, a gene associated to colon cancer metastasis [27]. Experimentally validated interactions are first identified computationally, thus, we expect to find at least one target site for miR-338-3p in MACC1 transcripts. But the interesting thing is that a second target site is segregating in human populations (Table 2). As multiple target sites are likely to increase the efficiency of the microRNA/gene interaction [11,28], the study of multiple segregating target sites may yield further insights in the variation at microRNA regulatory interactions in human populations. Importantly, high-throughput target experiments that do not rely on predictions using the reference genome will be useful in future studies.

**Table 2.**
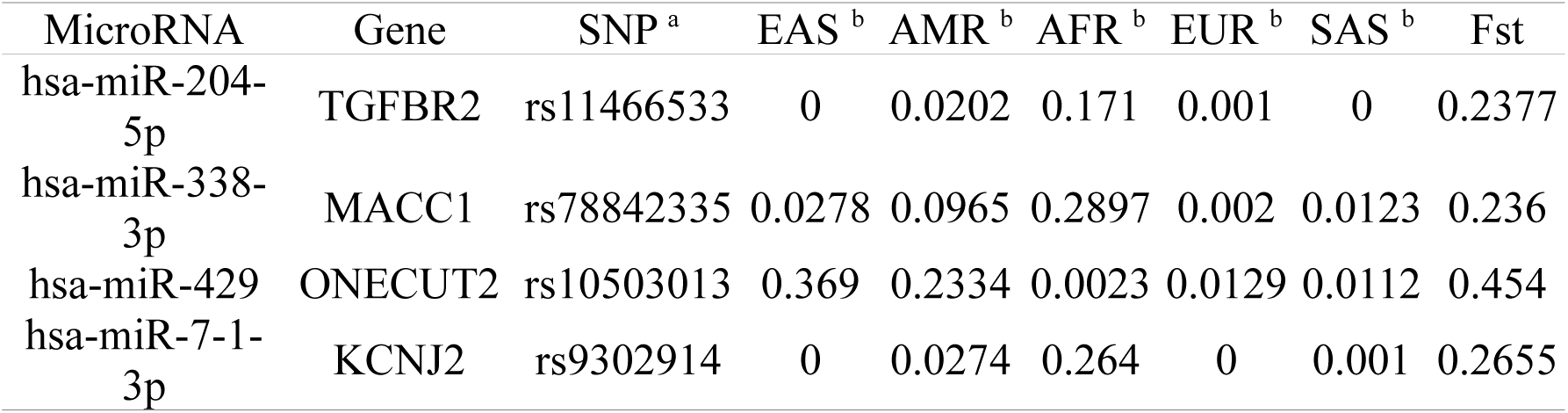
MicroRNA target sites with a very high differentiation among human populations.

In conclusion, here we show that the study of variation in human populations reveals the presence of microRNA target sites that are not detected with current methodological frameworks. We suggest that variation at target sites must be taken into account in biomedical studies as some populations (mostly of African origin) may be neglected. Further research is needed to quantify the impact of population variation is the function of microRNA target sites.

## Methods

Polymorphic sites at predicted microRNA target sites were retrieved from the PopTargs database (version 1; Hatlen, Helmy and Marco, under review; https://poptargs.essex.ac.uk). Only target sites for highly expressed microRNAs were considered. Population frequencies, Fst values and ancestral alleles were also retrieved in PopTargs, as provided in [29]. We only consider segregating alleles with a frequency between 0.01 and 0.99. Sequence alignment of primate sequences was extracted using the UCSC table browser [30]. The geographical location (latitude) of the genome samples sequenced in the 1000 genomes project was obtained from the sample descriptions at https://www.coriell.org/1/NHGRI/Collections/1000-Genomes-Collections/1000-Genomes-Project. When the specific location was ambiguous, we used the location of the capital city of the country of origin. For multiple collection points we computed the centroid middle point. Recently migrated populations were discarded (CHD, CEU, ASW, ACB, GIH, STU and ITU) were discarded from the analysis in Figure 2B. We extracted high-confidence experimentally validated microRNA targets from the miRTarBase database [31].

## Funding

This research was funded by Wellcome Trust, grant number 200585/Z/16/Z.

## Acknowledgments

The authors acknowledge the use of the High Performance Computing Facility (Ceres) and its associated support services at the University of Essex in the completion of this work.

